# DNA double-strand break signaling induces aberrant neuronal activity

**DOI:** 10.64898/2026.05.11.724110

**Authors:** Ping-Chieh Pao, Liwang Liu, L. Ashley Watson, Audrey Lee, Alexander Seguin, Daiyun Dong, Samaa Rasheed, Colin Staab, Oisín King, Ute Geigenmüller, Jay Penney, Fan Gao, Abdurrahman Muhtaseb, Ravikiran M. Raju, Li-Huei Tsai

## Abstract

Aberrant neuronal activity is an early pathological feature of numerous neurodegenerative disorders, including tauopathy, and is thought to play a role in disease progression. However, the mechanism underlying abnormal neuronal activity remains elusive. Here, we reveal a relationship between DNA double-strand break (DSB)/p53 pathway activation and aberrant neuronal activity. Activating p53 as part of the DNA damage response via DSB induction, or by preventing MDM2-mediated p53 degradation, causes aberrant activity in both mouse and human neurons. p53 activation induces the expression of genes regulating synaptic transmission, and p53-responsive gene upregulation is overrepresented in *postmortem* human Alzheimer’s disease neurons burdened with neurofibrillary tangles (NFTs). Using a human iPSC-based cerebral organoid model of frontotemporal dementia that exhibits relevant pathologies including elevated DSBs, aberrant neuronal activity, and NFTs, we show that inhibiting p53 transcriptional activity with a small molecule ameliorates aberrant calcium fluctuations in neurons. Together, our findings highlight p53 inhibition as a novel therapeutic strategy to counter aberrant neuronal activity in neurodegenerative diseases characterized by tauopathy.

## Introduction

Aberrant neuronal activity has been observed both in humans and rodents during neurodegeneration(Mucke & Selkoe, 2012; Palop & Mucke, 2016; Targa Dias Anastacio *et al*, 2022). Acute slices from biopsies of patients with early stage of Alzheimer’s disease (AD) exhibit increased neuronal bursts and concomitant upregulation of a gene signature linked to neuronal hyperactivity(Gazestani *et al*, 2023). Higher incident rates of new-onset seizures have also been observed in patients with AD, dementia with Lewy bodies, or frontotemporal dementia (FTD)(Beagle *et al*, 2017). In rodents, EEG and patch-clamp recordings revealed abnormal spiking in the J20 mouse model of amyloid pathology(Sanchez *et al*, 2012) and increased frequencies of spontaneous excitatory postsynaptic currents (sEPSCs) in mice expressing P301L mutant tau(Crimins *et al*, 2011), respectively. Emerging evidence also indicates that aberrant neuronal activity can itself exacerbate disease-associated pathology. Chemogenetically increasing neuronal activity accelerates tau pathology in mice expressing P301L mutant tau(Wu *et al*, 2016), and neuronal activity positively correlates with amyloid beta peptide production and amyloid deposition in mouse models(Bero *et al*, 2011; Yuan & Grutzendler, 2016). Moreover, AD patients with epileptiform activity show a faster decline in cognitive function(Vossel *et al*, 2016), and the antiseizure drug levetiracetam improves performance on spatial memory and executive function tasks in these patients(Vossel *et al*, 2021).

Emerging evidence from genomic studies highlights neuronal DNA damage in patients with neurodegeneration(Murdock & Tsai, 2023; Dileep *et al*, 2023). The single-cell whole-genome amplification assay revealed that neurons from patients with AD are particularly susceptible to DNA damage(Zhu *et al*, 2021), and transcriptional profiling of the AD prefrontal cortex (PFC) demonstrated an upregulation of genes associated with DNA damage and repair pathways, including the Cohesin-SA1/SA2 complex, *SIRT1, ATRX, USP47*, and *BAZ1B* (Mathys *et al*, 2023). *Postmortem* brain samples from patients with neurodegenerative disorders including mild cognitive impairment (MCI), AD, amyotrophic lateral sclerosis (ALS), and FTD show increased numbers of DSBs(Adamec *et al*, 1999; Wang *et al*, 2013; Shanbhag *et al*, 2019; Thadathil *et al*, 2021), and accrual of DSBs has also been found to correlate with cognitive decline in a healthy aging population study(Simpson *et al*, 2015). In addition, mutations in genes involved in DSB repair, such as *MRE11, NSB1, ATM*, and *LIG4*, can result in syndromes characterized by ataxia, microcephaly, and neurodegeneration(Madabhushi *et al*, 2014). Interestingly, exposing mice to DSB-inducing radiation led to increased synaptic transmission and upregulated levels of synaptic proteins(Marty *et al*, 2014; Rudobeck *et al*, 2017; Allen *et al*, 2020) as well as higher incident rates of secondary convulsive events in a rodent animal model of seizure(Jenrow *et al*, 2001). While these findings link the induction of DSBs to increased neuronal activity and hyperexcitability(Jenrow *et al*, 2001), the mechanism underlying this interaction remains poorly understood.

Here, we dissect the interaction between DSBs and aberrant neuronal activity using both mouse and human experimental systems. Utilizing patch-clamping techniques and multi-electrode array (MEA) recordings, we find that DSB induction through DSB repair blockade or site-specific endonuclease activation stimulates excitatory synaptic transmission, voltage-gated inward currents, and excitability in mouse cultured neurons. Transcriptomic analyses indicate that p53 activation stimulates the expression of genes associated with synaptic transmission and vesicle transport. Furthermore, the DSB/p53-dependent transcriptional response is associated with upregulation of synaptic transmission genes, which are overrepresented in *post-mortem* AD human excitatory neurons burdened with neurofibrillary tangles (NFTs).

We further confirm these findings in cerebral organoids derived from induced pluripotent stem cells (iPSCs) from an FTD patient carrying a heterozygous tau P301L mutation. The FTD organoids display classical mutant tau-associated pathology, including increased DSBs, tau hyperphosphorylation, and β-sheet conformational changes. Gene expression changes in neurons from the FTD organoids are associated with calcium ion pathways and upregulation of p53-target genes that are known to modulate synaptic transmission/function. Further, enriched pathway analysis indicates that genes upregulated both in FTD human neurons and in p53-activated mouse neurons have been associated with neuronal hyperexcitability phenotypes in humans. Finally, we show that FTD neurons exhibit activated DSB/p53 signaling and abnormal calcium fluctuations, which can be ameliorated by the blockade of p53 transcriptional activity with a small molecule. Together, our findings uncover a role for the DSB/p53 pathway in aberrant neuronal activity and highlight the importance of maintaining genome integrity to prevent neurodegeneration.

## Results

### Induction of DSBs facilitates excitatory synaptic transmission and leads to hyperexcitability in cultured mouse neurons

To examine the relationship between DSBs and neuronal activity, we induced DSBs in mouse cultured neurons at 15 days *in vitro* (DIV) using two different approaches and monitored neuronal electrophysiological properties by patch-clamp recordings. In the first approach, we blocked DSB repair with SCR7, a small molecule inhibitor of DNA ligase IV (Lig4i)(Srivastava *et al*, 2012). Lig4i treatment significantly increased DSB levels, as evidenced by upregulation of the comet tail moment and γH2AX immunostaining signal, but did not impact neuronal viability (**Figure 1a, b**). Patch-clamp recordings revealed a significant upregulation in spontaneous excitatory postsynaptic current (sEPSC) frequency in Lig4i-treated neurons (**Figure 1c**). Lig4i-treated neurons furthermore showed a higher frequency of action potentials in response to current stimulations, suggestive of hyperexcitability (**Figure 1d**). Given that voltage-gated sodium channels are key players in action potential initiation in neurons, we next measured the inward current after Lig4i treatment. Compared to solvent-treated neurons, Lig4i-treated neurons exhibited high inward currents upon application of voltage (**Figure 1d**). These findings indicate that Lig4i treatment stimulates excitatory synaptic transmission, voltage-gated inward current, and excitability in cultured neurons.

**Figure 1.**
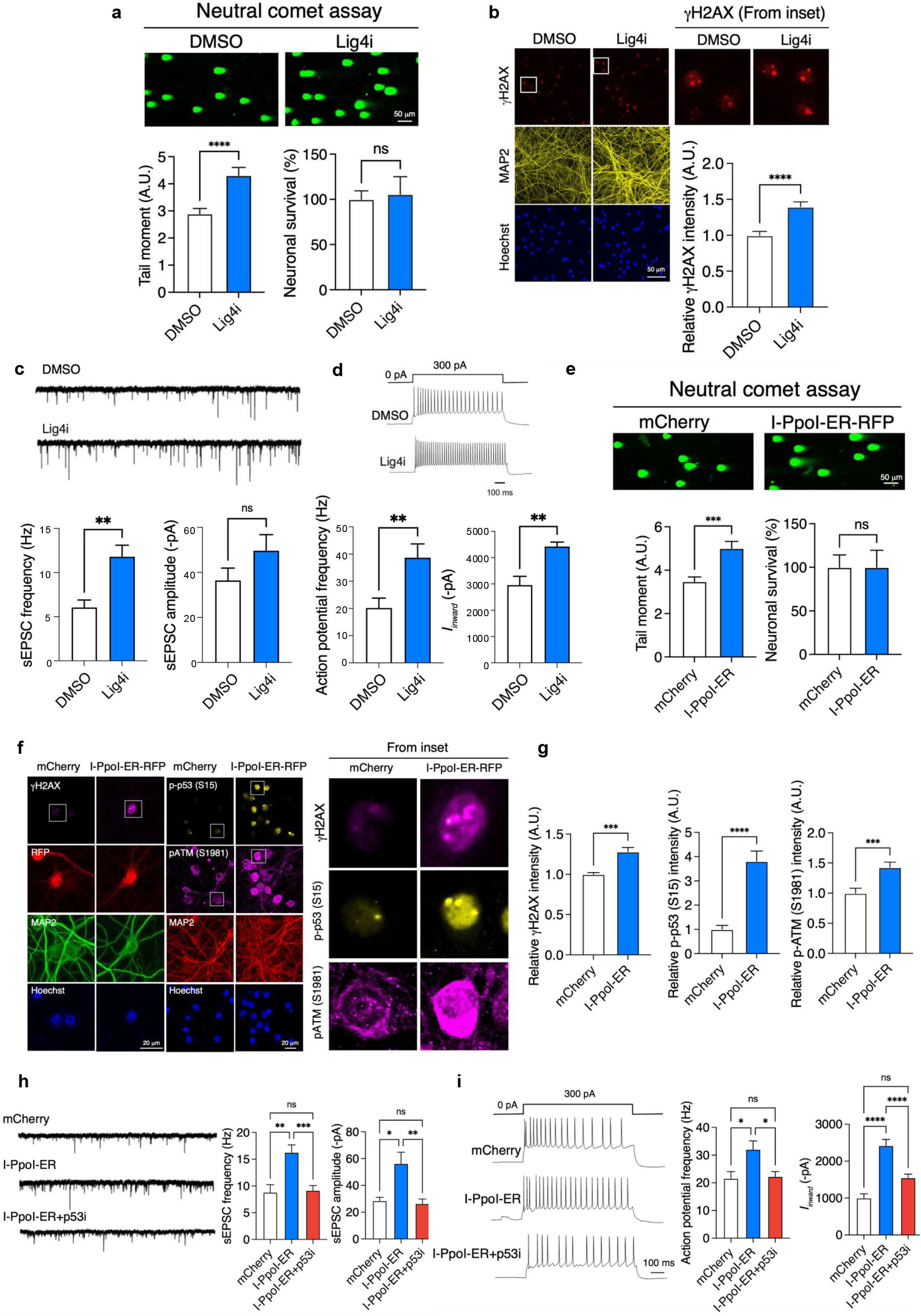
DSBs induce aberrant neuronal activity and hyperexcitability through the activation of p53. **a** Representative images and quantification of the neutral comet assay using cultured neurons treated with Lig4i or DMSO control (n=243, 279). Neuronal viability assay includes 14 replicates per group. **b** Representative images of cultured neurons stained for γH2AX (red), MAP2 (yellow), and Hoechst (blue). Bar graph represents the intensity of γH2AX from 3 images per group (n=90, 84). **c** Representative sweeps of sEPSC from DMSO and Lig4i-treated groups. Bar graphs represent sEPSC frequency and amplitude (n=21, 27). **d** Representative sweeps showing voltage responses from DMSO and Lig4i-treated groups elicited by 800 ms current injections of 300 pA. Bar graphs represent action potential frequency (n=22, 26) and inward current (n=13, 15). **e** Representative images and quantification of the neutral comet assay using cultured neurons expressing I-PpoI-ER-RFP or mCherry control (n=291, 386). Neuronal viability assay includes 16 replicates per group. **f** Left, representative images of cultured neurons stained for γH2AX (magenta), MAP2 (green), and Hoechst (blue). Right, representative images of cultured neurons stained for p-p53 (S15) (yellow), p-ATM S1981 (magenta), MAP2 (red), and Hoechst (blue). Bar graphs represent the intensity of γH2AX (n=9, 9) and p-p53 (S15) and p-ATM (S1981) (n=69, 74). **h** Representative sweeps of sEPSC from neurons expressing mCherry, I-PpoI-ER, and I-PpoI-ER treated with p53 inhibitor (Pifithrin-α; p53i). Bar graphs represent sEPSC frequency and amplitude (n=14, 27, 18). **i** Representative sweeps showing voltage responses elicited by 800 ms current injections of 300 pA. Bar graphs represent action potential frequency (n=22, 40, 30) and inward current (n=14, 33, 37). All values are shown as mean ± SEM. Statistical analysis: **a-e, g** two-tailed Student’s t-test; **h, i** one-way ANOVA with Tukey’s post hoc test. Source data are provided as a Source Data file.

In our second approach, we induced DSBs by transducing the neurons with lentivirus expressing I-PpoI, a rare-cutting endonuclease that generates DSBs at defined genomic loci once it enters the nucleus. By fusing I-PpoI to the estrogen receptor, the endonuclease is trapped in the cytoplasm until addition of 4-hydroxytamoxifen (4-OHT), which binds to the estrogen-receptor domain, triggering nuclear translocation of I-PpoI(Dobbin *et al*, 2013). As expected, the addition of 4-OHT in I-PpoI-ER-expressing neurons led to an increase in DSBs as evidenced by a larger comet tail moment (**Figure 1e**) and higher γH2AX signal (**Figure 1f, g**). Patch-clamp recordings showed that activation of I-PpoI led to similar electrophysiological changes as were seen in Lig4i-treated neurons, including increased frequency/amplitude of sEPSC, upregulated inward current, and hyperexcitability (**Figure 1h, i**). Thus, our findings show that induction of DSBs triggers hyperactivity and hyperexcitability in cultured neurons.

Since the formation of DSBs is known to activate the DNA damage response (DDR) pathway(Madabhushi *et al*, 2014), we investigated how modulation of the DDR impacted the effect of DSB induction on neuronal activity. Compared to controls, neurons with activated I-PpoI indeed exhibited higher intensity of p-p53 S15 and pATM S1981 staining (**Figure 1f, g**), indicative of DDR pathway activation. To modulate the DDR, we focused on p53, an important effector at the hub of the DDR pathway. Inhibiting p53 transcriptional activity with Pifithrin-α (p53i) while inducing DSBs with I-PpoI activation significantly ameliorated aberrant synaptic transmission, hyperexcitability, and aberrant inward current in neurons (**Figure 1h, i**), suggesting that p53 activation plays a role in mediating the effect of DSBs on neuronal hyperactivity. Consistently, pharmacological inhibition of p53 with Pifithrin-α also reduced aberrant synaptic transmission and hyperexcitability in neurons treated with Lig4i (**Supplementary Figure 1a–e**). In contrast, resting membrane potential and input resistance, two determinants of neuronal excitability, remained unchanged across these conditions (**Supplementary Figure 1f, g**).

### Activation of p53 signaling triggers aberrant neuronal activity and upregulates the expression of genes linked to synaptic transmission

To further examine the role of p53 activation in neuronal activity and synaptic transmission, we treated cultured neurons with Nutlin-3. Nutlin-3 is a small molecule known to augment p53 levels by preventing MDM2-mediated p53 degradation(Vassilev *et al*, 2004). We found that Nutlin-3 increased the frequency/amplitude of sEPSC, promoted high inward currents, and led to hyperexcitability in cultured neurons (**Figure 2a, b**) with no change in resting membrane potential and input resistance (**Supplementary Figure 1h, i**). Results from patch-clamp recording were confirmed by similar observations using multi-electrode array (MEA) recordings, which showed that treatment with Nutlin-3 increased the spontaneous activity in cultured neurons (**Figure 2c**). Together, our findings suggest that DSBs trigger aberrant neuronal activity through activation of p53.

**Figure 2.**
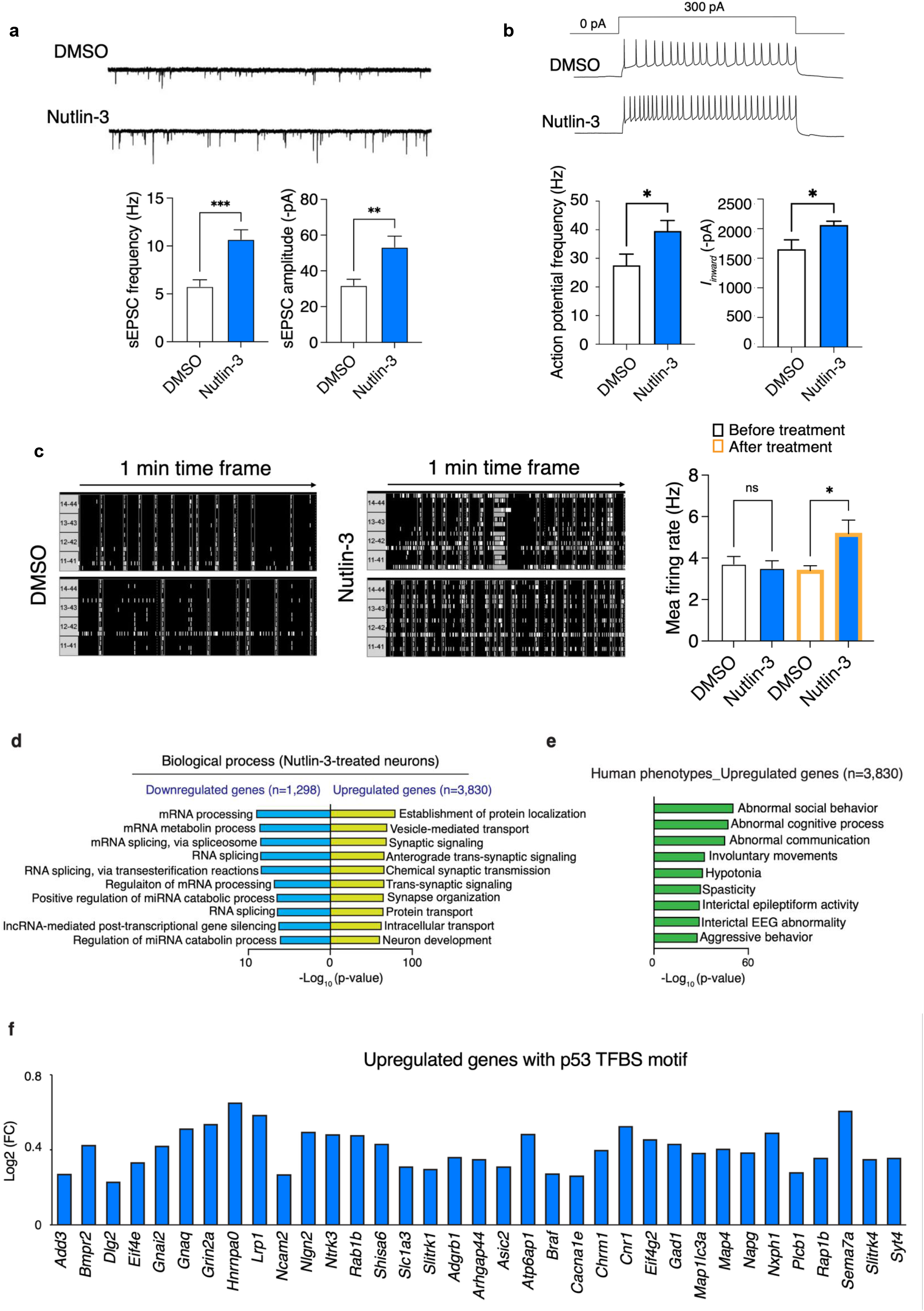
Activation of p53 signaling upregulates expression of genes linked to synaptic transmission. **a** Representative sweeps of sEPSC from neurons treated with p53 activator (Nutlin-3) or DMSO control. Bar graphs represent sEPSC frequency and amplitude. (n=19, 24). **B** Representative sweeps showing voltage responses elicited by 800 ms current injections of 300 pA. Bar graphs represent action potential frequency (n=12, 11) and inward current (n=19, 24). **c** Representative raster plots of MEA recordings showing spontaneous activity. Bar graphs represent the mean firing rate of cultured neurons before and after the treatment of the p53 activator (Nutlin-3) (n=16, 16). GO terms (**d**) and disease terms (**e**) enriched for DEGs were annotated using Toppgene. **f** Bar graph showing the Log_2_(Fold change) of selected genes that are predicted to have p53 motif in the promoter regions. p53 target genes were extracted from the transcription factor binding site (TFBS) database (V$P53_02 and V$P53 DECAMER Q2). All values are shown as mean ± SEM. Statistical analysis: **a, b** two-tailed Student’s t-test; **c** one-way ANOVA with Tukey’s post hoc test. Source data are provided as a Source Data file.

To understand how p53 promotes neuronal activity, we performed bulk RNA-sequencing (RNA-seq) of cultured neurons treated with Nutlin-3 or DMSO as a solvent control. We identified 1,298 downregulated and 3,830 upregulated genes in Nutlin-3 treated neurons relative to controls (*p*.*adj* <0.05). Downregulated genes were involved in mRNA processing and splicing pathways, while upregulated genes were enriched for gene ontology terms related to synaptic transmission/signaling and vesicle transport (**Figure 2d**).

We next focused on genes that are upregulated after p53 activation. We found that genes upregulated in response to p53 activation overlapped with gene sets associated with abnormal cognition, EEG abnormality, epileptiform activity, and seizure phenotypes in humans (**Figure 2e**). Further, upregulated but not downregulated genes were statistically enriched for p53 target genes (88 upregulated and only 5 downregulated genes were found among the total 264 expressed p53 target genes; hypergeometric test; p-value: 1.36E-05 and 0.999988, respectively) and many of the p53 target upregulated genes are also known to modulate synaptic transmission/function and excitability, including *Grin2a, Slc4a4, Slc1a3, Dlg2, Ncam2*, and *Cacna1e* (**Figure 2f**). These observations suggest that p53 activation stimulates the expression of genes that promote synaptic transmission and are associated with epileptic-like phenotypes.

### *Postmortem* AD human neurons burdened with neurofibrillary tangles show gene expression signatures of p53 activation and elevated DSB levels

Recent genomic studies demonstrated increased expression of genes related to the DNA damage pathway in neurons from patients with AD(Zhu *et al*, 2021; Mathys *et al*, 2023; Dileep *et al*, 2023). We therefore sought to determine whether gene signatures of p53 activation could be detected in human AD neurons. We first compared the upregulated genes (UP_DEGs) observed here in cultured mouse neurons treated with the p53 activator Nutlin-3 (p53-responsive gene signature) to UP_DEGs previously reported in two different datasets for human excitatory (Ex) neurons from *postmortem* AD brains(Otero-Garcia *et al*, 2022; Mathys *et al*, 2023). Interestingly, we observed significant overlap of Nutlin-3-responsive UP_DEGs with genes found to be upregulated in human AD excitatory neurons in both datasets (**Figure 3a-b**). Overlap with UP_DEGs in excitatory neurons associated with AD global pathology was especially pronounced in deep cortical layers (**Figure 3a**, Ex L6 CT cluster; hypergeometric test; p-value: 1.62E-128; overlapping genes=492). Further, we found a striking degree of overlap between Nutlin-3 upregulated genes and UP_DEGs in excitatory neurons bearing neurofibrillary tangles (NFTs) (**Figure 3b**; Ex1, Ex2, Ex7). Notably, upregulated genes that overlap with cluster Ex2, one of the most p53-response enriched clusters (**Fig. 3c**, Ex2, hypergeometric test; *p*-value: 2.11E-243; overlapping genes=493), are linked to synaptic transmission/seizure phenotypes (**Figure 3d**). These findings indicate that human AD neurons, particularly those with NFTs, exhibit a similar transcriptional signature to that observed following p53 activation in mouse cultured neurons.

**Figure 3.**
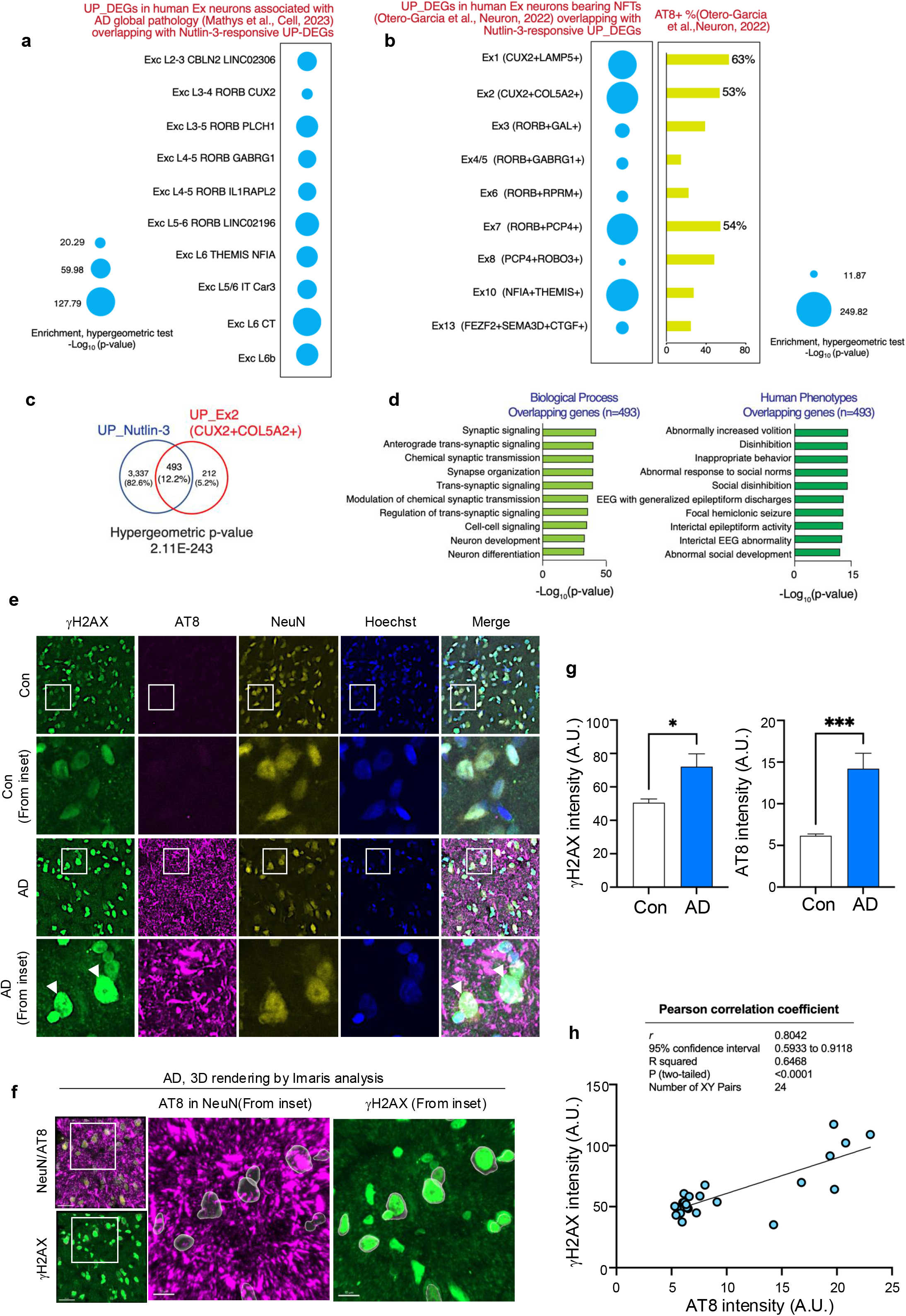
A p53-associated synaptic gene upregulation is conserved in AD human neurons with neurofibrillary tangles (NFTs) **a** The plot shows the overlap between UP_DEGs in p53 activated cultured mouse neurons and UP_DEGs in AD human neurons(Mathys *et al*, 2023). The size of bubble represents the statistically significance of overlap between two gene lists. **b** (Left) The plot shows the overlap between DEGs in p53 activated cultured mouse neurons and DEGs in AD human neurons with NFT(Otero-Garcia *et al*, 2022). The size of bubble represents the statistically significance of overlap between two gene lists. (Right) The plot shows the percentage of NFT-bearing neuron in each cluster. **c** Venn diagram representing the overlap of DEGs between groups. **d** GO terms and disease terms enriched for overlapping upregulated genes were annotated using Toppgene. **e** Representative images of post-mortem human brain from health subjects (n=3) or patients with AD (n=3). Brain sections were stained for NeuN (yellow), γH2AX (green), AT8 (magenta), and Hoechst (blue). Arrowhead indicate AD neurons contains high levels of γH2AX and AT8 signals. Four sections per subject were analyzed. **f** 3D rendering analysis using Imaris software. NeuN signals were used to create masks for quantifying intensity of γH2AX and AT8 in AD neurons. **g** Bar graphs represent γH2AX and AT8 intensity in control and AD group. **h** Plot shows the linear correlation between mean intensity of γH2AX and mean intensity of AT8 within NeuN surface created by 3D rendering analysis. All values are shown as mean ± SEM. Statistical analysis: **a-c** hypergeometric test. **g** two-tailed Student’s t-test **h** Pearson correlation coefficient. Source data are provided as a Source Data file.

Formation of NFTs is a defining characteristic of tauopathy, which is also accompanied by increased DSB levels and aberrant neuronal activity(Asada-Utsugi *et al*, 2022). In fact, inducing DSBs with the genotoxin etoposide promotes tau pathology in mouse neurons(Asada-Utsugi *et al*, 2022). To validate the presence of elevated DSBs in human NFT-bearing neurons, we performed immunolabeling for γH2AX, AT8, and NeuN in *postmortem* prefrontal cortex (PFC) tissue from individuals with AD and non-AD controls. Three-dimensional rendering analysis revealed increased γH2AX and AT8 signal intensities within NeuN surfaces in the AD group compared to the non-AD group. Furthermore, γH2AX intensity positively correlated with AT8 intensity within NeuN surfaces in AD samples (Pearson correlation coefficient, *r*= 0.8042, *p-*value < 0.0001) (**Figure 3e–h**). Together, these findings suggest that tau pathology in the human brain is positively correlated with DSBs and a p53-dependent transcriptional signature that includes upregulation of synaptic transmission genes.

### Activation of the DSB/p53 signaling pathway promotes neuronal calcium events in cerebral organoids derived from a healthy human control iPSC line

We next tested if the observed effects of DSB induction or direct p53 activation on neuronal activity could be replicated in cerebral organoids derived from a healthy human control iPSC line (CS-0009-01). Immunostaining analysis demonstrated that the cerebral organoids generated from these iPSCs (**Figure 4a)** robustly expressed pan-neuronal markers (NeuN, MAP2) as well as markers of cortical neurons (CTIP2, TBR1) and synapses (SVP38) (**Figure 4b; Supplementary Figure 2**).

**Figure 4.**
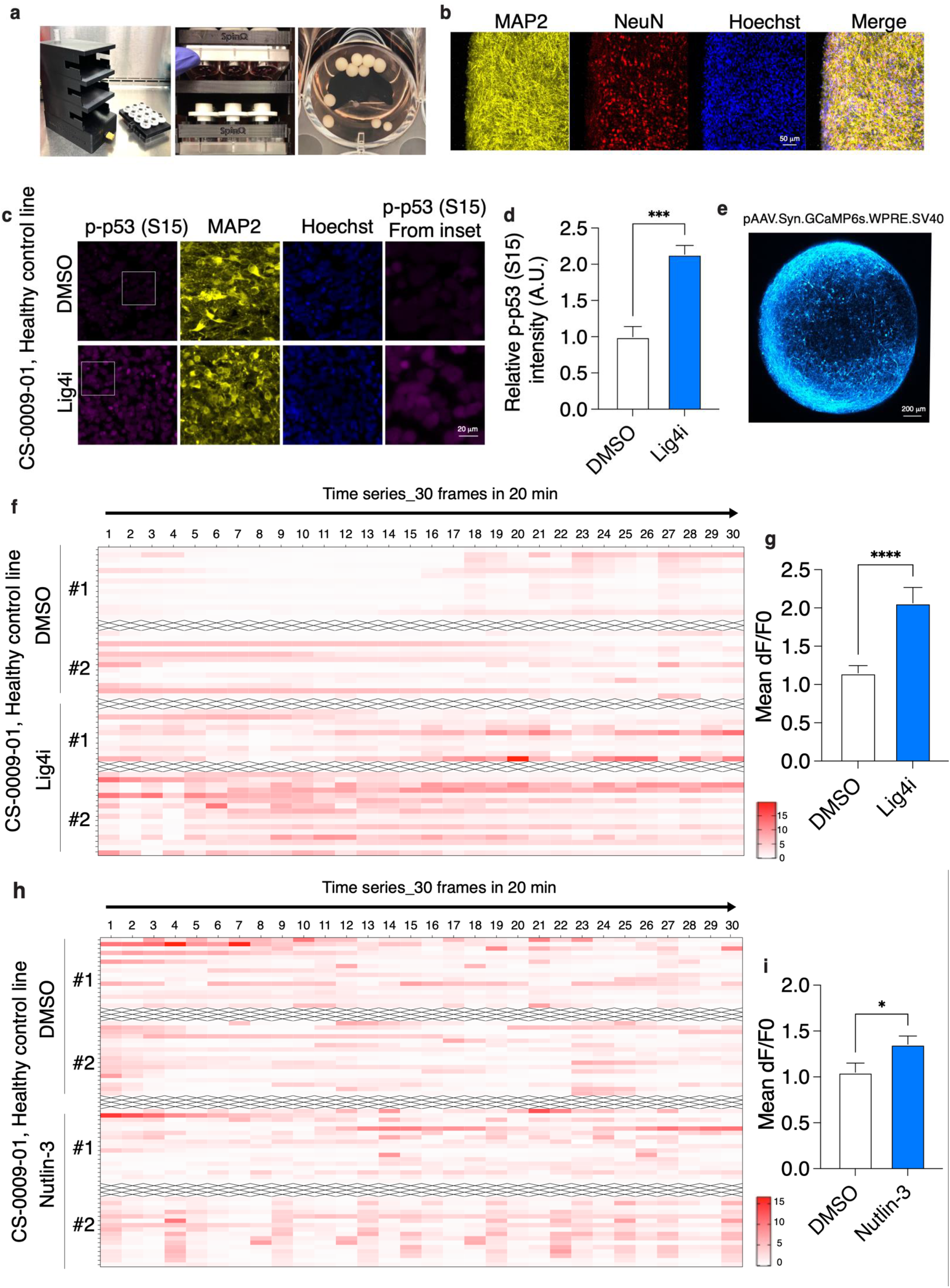
Activation of DSB/p53 pathway stimulates neuronal firing in 3D cerebral organoids derived from a healthy iPSC line. **a** Pictures of cerebral organoids derived from a healthy iPSC line (CS-0009-01) and 3D-printed bioreactor system. **b** Representative images of 2-month-old cerebral organoids stained for markers for pan-neuronal markers (β3-tubulin, MAP2, NeuN). **c,d** Representative images of 4-month-old cerebral organoid sections stained for p-p53 S15 (magenta) and Hoechst (blue). Two images per organoid and 3 organoids per group examined. Bar graph represents the immunoreactivity for p-p53 S15 normalized to DMSO treatment. Three organoids and two sections per organoid were analyzed in each group. **e** Representative images of a 5-month-old cerebral organoid transduced with pAAV.Syn.GCaMP6s.WPRE.SV40 for calcium imaging. **f** Heatmap representing the calcium events (deltaF/F0) in a time series video. **g** Bar graph represents the mean deltaF/F0 in a time series video. (n=74, 58). Four organoids per group. **h** Heatmap representing the calcium events (deltaF/F0) in a time series video. **i** Bar graph represents the mean deltaF/F0 in a time series video. **i** (n=59, 96). At least 5 organoids per group were analyzed. All values are shown as mean ± SEM. Statistical analysis: **d, g, i** two-tailed Student’s t-test. Source data are provided as a Source Data file.

We first determined the effects of indirectly activating p53 by inducing DSBs in the cerebral organoids. As expected, organoids treated with Lig4i to block DNA repair exhibited p53 activation, as demonstrated by an increase in p53 phosphorylation at S15 (**Figure 4c, d**). To examine neuronal activity in these organoids, we used calcium imaging. Organoids were transduced with adeno-associated virus (AAV) expressing the genetically encoded calcium indicator GCaMP6 under the control of a neuronal-specific *SYN1* promoter (**Figure 4e**). Live fluorescence imaging of Lig4i-treated organoids showed elevated amplitudes for calcium fluctuation (ΔF/F0) compared to vehicle-treated organoids (1.79-fold; **Figure 4f, g**), indicating that DSB induction led to increased neuronal calcium events. We then used Nutlin-3 to directly activate the p53 pathway in the cerebral organoids and again observed increases in the amplitude of ΔF/F0 (1.29-fold; **Figure 4h, i**). These findings suggest that activation of the DSB/p53 pathway stimulates neuronal activity in human cerebral organoids.

### Aberrant neuronal calcium events in an iPSC model of tauopathy is ameliorated by p53 inhibition

To further investigate the causal relationship between DSBs, p53 activation, and neuronal hyperactivity in the context of neurodegeneration, we constructed a cerebral organoid model of tauopathy. To this end, we used iPSCs from an FTD patient heterozygous for the P301L-coding mutation in *MAPT*(Seo *et al*, 2017), as well as two control isogenic iPSC lines where the *MAPT* mutation had been corrected by CRISPR/Cas9 editing (**Figure 5a, Supplementary Figure 3a**). Cerebral organoids generated from the FTD line recapitulated various mutant tau-associated pathologies. Specifically, 60-day-old FTD organoids showed increased levels of hyperphosphorylated tau (p-tau) by Western blot analysis (T181, S396, and S202; **Supplementary Figure 3b, c**), with a pronounced p-tau increase in the cell soma (T181, S396, S404, and S202/T205; **Figure 5b, c**) that is reminiscent of the somatodendritic accumulation of p-tau observed in humans with AD(Merino-Serrais *et al*, 2013). Staining of FTD organoids with thioflavin S further revealed the increased β-sheet conformations characteristic of aggregated tau (**Supplementary Figure 3d, e**). FTD organoids also displayed elevated levels of neuronal DSBs compared to control organoids, as demonstrated by elevated γH2AX foci detected by IHC (**Figure 5d, e**) and a higher tail moment from FACS isolated NeuN+ nuclei in the comet assay (**Figure 5f, g, Supplementary Figure 3f**). These findings suggest that FTD organoids represent a faithful model of human tauopathy.

**Figure 5.**
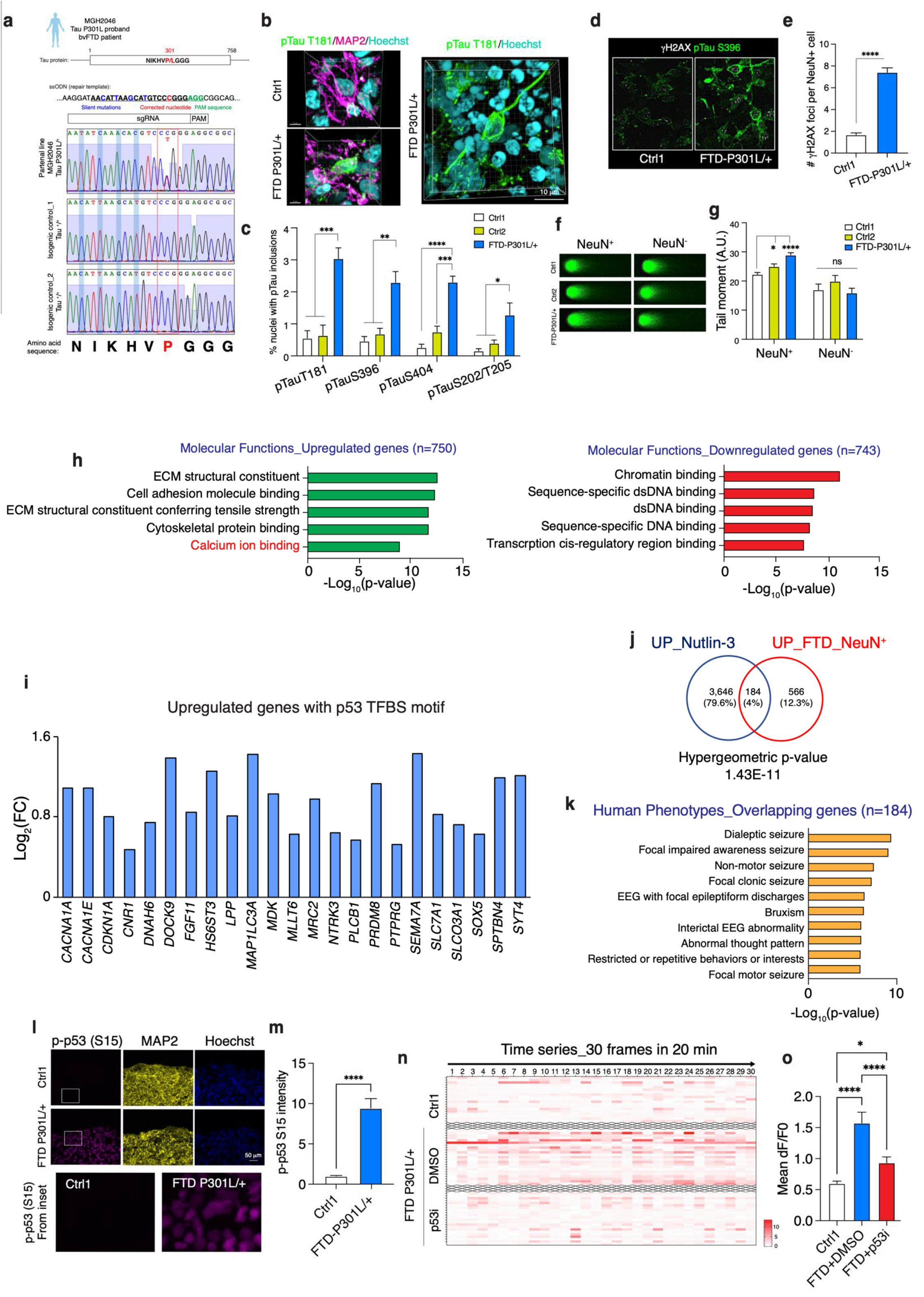
FTD cerebral organoids recapitulate tauopathy, exhibit activated DSB/p53 pathway activation and aberrant firing. **a** Schematics of isogenic controls generated from an FTD patient iPSC line carrying a heterozygous Tau P301L mutation. Sanger sequencing confirms the insertion of a desired repair template into an FTD patient iPSCs. **b** 3D rendering representative images of somatodendritic accumulation of ptau T181 (green), MAP2 (magenta), and Hoechst (cyan) in 60-day-old organoids. **c** Bar graphs represent the percentage of nuclei with ptau (T181, S396, S404, and S202/T205) somatodendritic inclusion. **d** Representative images of organoid sections stained for pTau S396 (green), γH2AX (white), NeuN (magenta), and Hoechst (cyan). **e** Bar graph represents the number of γH2AX foci in NeuN^+^ nuclei. **f** Representative images of neutral comet assay on NeuN^+^ nuclei isolated from 90-day-old organoids. **g** Bar graph represents the tail moment in NeuN^+^ and NeuN-nuclei isolated from organoids. **h** GO terms enriched in DEGs were annotated using Toppgene. **i** Bar graph showing the Log_2_(Fold change) of upregulated genes that predicted to have p53 motif in the promoter regions. **j** Venn diagram representing the overlap of DEGs between groups. **k** Human disease terms enriched in overlapping DEGs were annotated using Toppgene. **l** Representative images of p-p53 S15 (magenta), MAP2 (yellow), and Hoechst (cyan) in organoids. **m** Bar graph represents the intensity of p-p53 S15 in organoids. Three organoids were examined. **n** Heatmap representing the calcium events (deltaF/F0) in a time series video. **o** Bar graph represents the mean deltaF/F0. (n=111, 47, 68 from at least 3 organoids per group). All values are shown as mean ± SEM. Statistical analysis: **e, m** two-tailed Student’s t-test; **c, g, o** one-way ANOVA with Tukey’s post hoc test.; **j** hypergeometric tests. Source data are provided as a Source Data file.

We then determined gene expression changes in neurons within the FTD organoid model. Transcriptomic analysis of NeuN^+^ nuclei isolated from 90-day-old FTD and control organoids (**Supplementary Figure 4a**) identified 750 upregulated and 743 downregulated genes in FTD organoids relative to controls (*FDR*<0.05; **Supplementary Figure 4b**). Upregulated genes were involved in the regulation of extracellular matrix (ECM), cell adhesion, and calcium pathways, whereas the downregulated genes were more specific to DNA binding (**Figure 5h**). A subset of the upregulated genes contains a consensus p53 binding motif and are known to modulate synaptic transmission/function, including *CACNA1A, CACNA1E, PLCB1, PTPRG, SEMA7A, and SYT4* (**Figure 5i**). We further found that 184 of the 750 FTD-upregulated genes were also upregulated by p53 activation in cultured mouse neurons (hypergeometric test, *p*=1.43E-11, **Figure 5j**). Importantly, these overlapping upregulated genes were associated with seizure phenotypes in human GO analysis (**Figure 5k**). The gene expression analysis of the FTD organoids suggests an association of p53 activation and increased neuronal activity.

To experimentally validate an association of p53 activation and increased neuronal activity in FTD organoids, we examined serine 15 phosphorylation of p53, a modification that occurs downstream of DNA damage to stabilize p53(Lavin & Gueven, 2006) (**Figure 4**), and measured neuronal calcium dynamics, respectively. FTD organoids showed both p53 activation (pS15) (**Figure 5l, m**) and increased calcium fluctuations – indicative of enhanced neuronal activity – compared to isogenic control organoids (2.62-fold; **Figure 5n, o**). Importantly, inhibition of p53 with Pifithrin-α attenuated the enhanced neuronal activity (**Figure 5n, o**), directly linking p53 activation to aberrant neuronal activity in this FTD disease model. Together, our observations suggest that DSBs in FTD patient-derived neurons lead to p53 activation, which in turn promotes the expression of genes that enhance synaptic transmission and aberrant neuronal activity. Our findings highlight p53 inhibition as a novel therapeutic strategy to counter aberrant neuronal activity in neurodegenerative diseases characterized by tauopathy.

## Discussion

This study uncovers a causal relationship between the DSB/p53 signaling pathway and aberrant neuronal activity. Activation of the p53 pathway—either through DSB induction or by using Nutlin-3 to prevent MDM2-mediated p53 degradation—leads to abnormal activity in both mouse and human neuron cultures. In a human iPSC-derived cerebral organoid model of FTD, which recapitulates many tau-associated pathologies including active DSB/p53 signaling and abnormal neuronal activity, inhibiting p53 transcriptional activity with Pifithrin-α (p53i) reduces abnormal calcium fluctuations in neurons. These findings indicate that p53 plays a key role in aberrant neuronal activity observed in neurons burdened with DSBs in disease conditions such as tauopathies.

We have previously reported that Topoisomeriase 2B-mediated DSBs in the promoters of immediate early genes (IEGs) are essential for IEG expression upon neuronal activity stimulation(Madabhushi *et al*, 2015). Here, we show a relationship between DSBs and aberrant neuronal calcium fluctuations in pathological conditions. These observations indicate that an excess of DSBs could drive aberrant neuronal activity and that sufficient DSB repair capacity is critical to maintain proper brain function. In this study, we applied two different approaches to induce DSBs – blocking DSB repair and endonucleolytic DSB generation at defined genomic loci – that are likely to generate DSBs at different genomic locations.

The observation that both approaches led to similar electrophysiological consequences implies that downstream DNA damage signaling rather than the specific locations of the DNA breaks might be crucial for stimulating aberrant neuronal activity. Sustained activation of I-PpoI may recruit and sequester DSB repair machinery to I-PpoI cutting sites, thereby reducing the cell’s ability to repair physiological DSBs, and DNA ligase 4 inhibition likely hinders repair of physiological DSBs in a more direct manner. Regardless of the exact locations of DSBs, both approaches to induce genomic damage led to activation of p53 and altered expression of genes that are predicted to impact neuronal activity.

p53 is a transcription factor whose targets have been shown to include many synaptic genes. Genome-wide p53 ChIP-chip analysis in Drosophila neurons revealed enrichment of p53 at promoters of genes that regulate synaptic transmission/vesicles, and p53 knockdown was associated with reduced levels of synaptic genes in mouse cultured neurons(Merlo *et al*, 2014). We observed similar p53-dependent gene expression changes here, as well as identifying a gene signature associated with synaptic transmission/function that is shared among genes upregulated by p53 activation or by tauopathy in neurons. These findings suggest that the aberrant neuronal activity observed in tauopathy could be, in part, a result of the p53-mediated transactivation of genes modulating synaptic transmission/functions.

Our findings show that blocking p53 activation attenuates aberrant activity in FTD cerebral organoids. A similar protective effect of p53 inhibition has also been demonstrated in a Rett Syndrome brain organoid model(Samarasinghe *et al*, 2021). Given that increased DSB levels and aberrant neuronal activity have also been reported in neurodegenerative disorders other than tauopathies, p53 activation may represent a common pathogenic mechanism shared among brain diseases. In addition, other DNA repair factors may be involved in driving aberrant neuronal activity. Previous studies have indicated that reduced levels of BRCA1 in neurons are associated with hyperexcitability(Suberbielle *et al*, 2015), and emerging evidence indicates a role for single-strand breaks and PARP1 hyperactivity in triggering aberrant neuronal activity(Komulainen *et al*, 2021). These observations suggest that aberrant neuronal activity might be modulated by multiple pathways that are activated in response to genomic instability. As the present study primarily focuses on excitatory neurons, future work will be needed to determine whether DNA damage also disrupts inhibitory neuronal function and thereby contributes to excitation–inhibition (E–I) imbalance.

Aberrant neuronal activity has been shown to modulate disease pathogenesis in several studies: inducing neuronal activity is associated with increased amyloid-β peptide generation and augmented plaque deposition in amyloid mouse models (Bero *et al*, 2011; Yuan & Grutzendler, 2016), augmented aggregation/spreading of α-synuclein in mice(Wu *et al*, 2020), and increased growth and spread of tumor cells in diffuse gliomas and glioblastoma(Venkatesh *et al*, 2015). A recent genomic study also showed that gene fusion events are overrepresented in excitatory neurons with severe NFT pathology and DNA damage in AD patients(Dileep *et al*, 2023), suggesting that DNA damage could perturb neuronal function via multiple mechanisms. Therefore, boosting DSB repair or blocking p53 activation in neurological diseases with DSB accumulation and aberrant neuronal activity would likely provide beneficial effects and help to attenuate pathogenesis. However, given the critical tumor suppressor function of p53, its inhibition may also pose a potential risk of facilitating tumor development or progression.

In summary, our work establishes a causal link between the DSB/p53 pathway and aberrant neuronal activity observed in tauopathies, with potential relevance to other neurodegenerative disorders. Our study underscores the importance of maintaining genome integrity to prevent neurodegeneration, suggesting the DSB repair machinery as a potential therapeutic target for the treatment of neurodegenerative disorders.

## Supplementary Figure legends

**Supplementary Figure 1.**
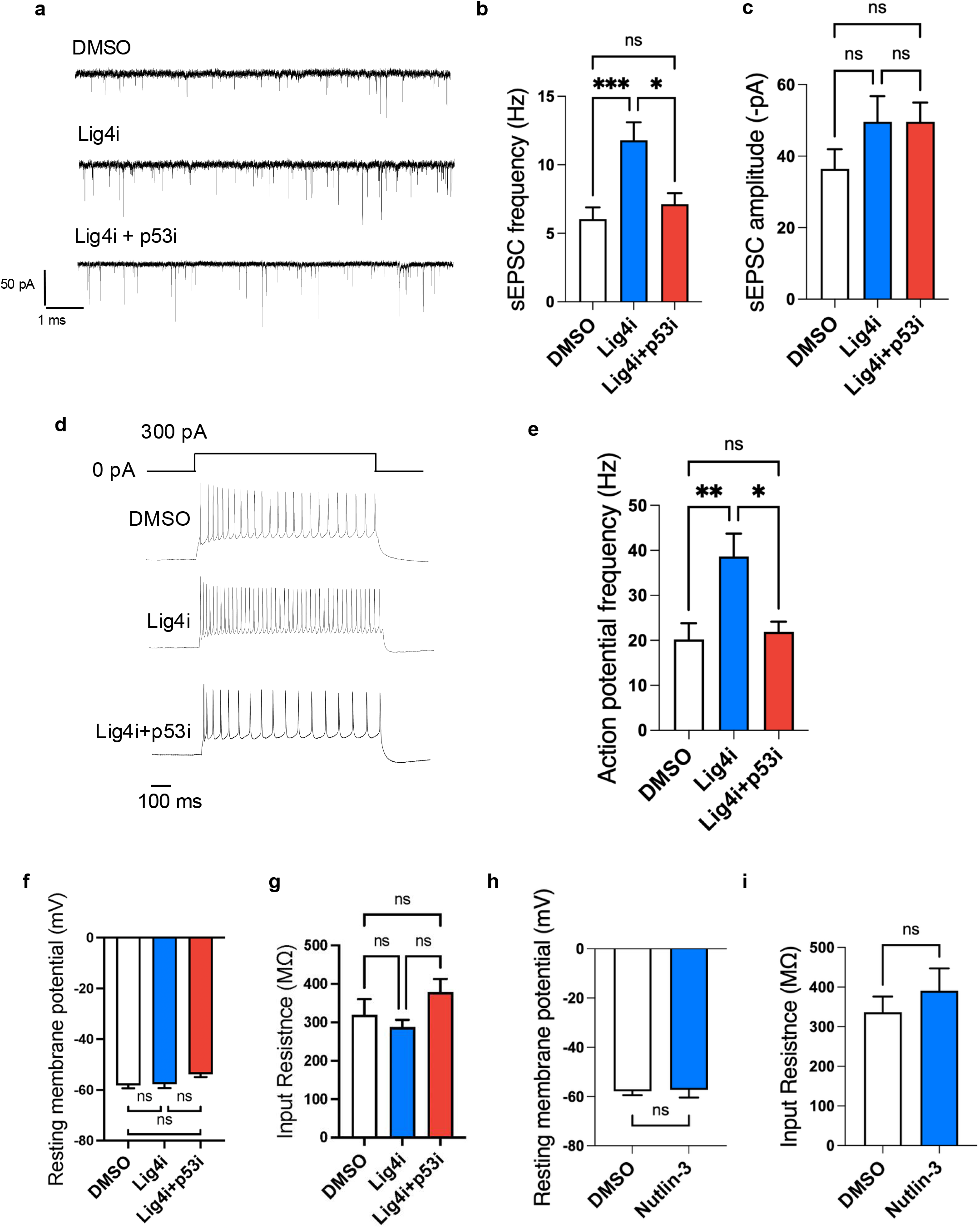
Electrophysiological characterization of mouse culture neurons burdened with DSBs.1. **a-c** Representative sweeps of sEPSC from neurons expressing DMSO, Lig4i, and Lig4i treated with p53 inhibitor (Pifithrin-α; p53i). Bar graphs represent sEPSC frequency and amplitude (n=21, 27, 18). **d-g** Representative sweeps showing voltage responses elicited by 800 ms current injections of 300 pA. Bar graphs represent action potential frequency (n=22, 26, 19), resting membrane potential (n=24, 25, 23) and input resistance (n=15, 25, 22). **h-i** Bar graphs represent resting membrane potential (n=14, 13) and input resistance (n=12, 11) in neurons treated with Nutlin-3 or DMSO vehicle control. All values are shown as mean ± SEM. Statistical analysis: **b,c,e-g** one-way ANOVA with Tukey’s post hoc test, **h,i** two-tailed Student’s t-test; Source data are provided as a Source Data file.

**Supplementary Figure 2.**
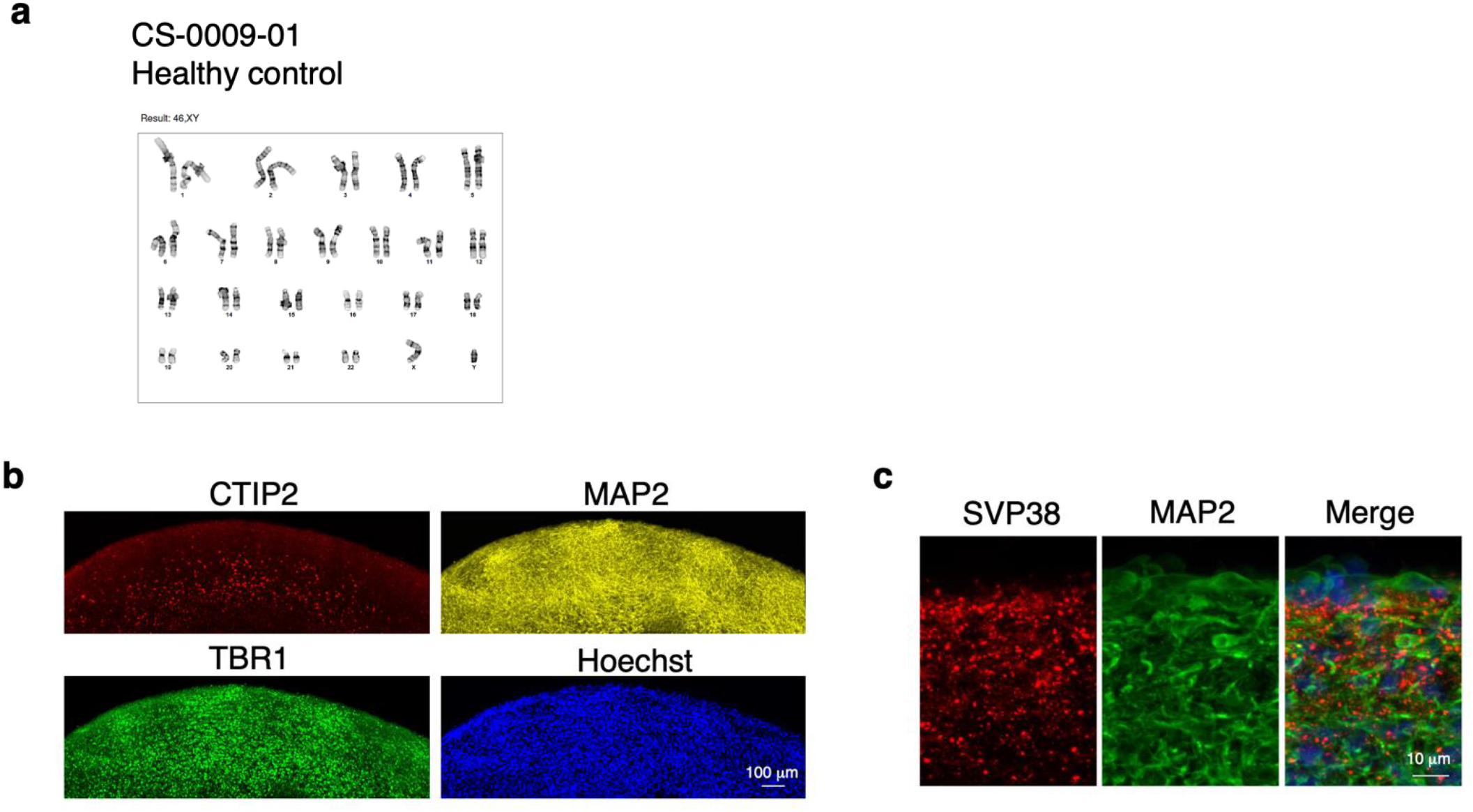
Characterization of human cerebral organoids derived from healthy iPSC line (CS-0009-01 line) **a** Normal karyotype of CS-0009-01 iPS line. **b,c** Representative images of 2-month-old cerebral organoids stained for markers for cortical neurons (CTIP2, TBR1) and synapse (SVP38).

**Supplementary Figure 3.**
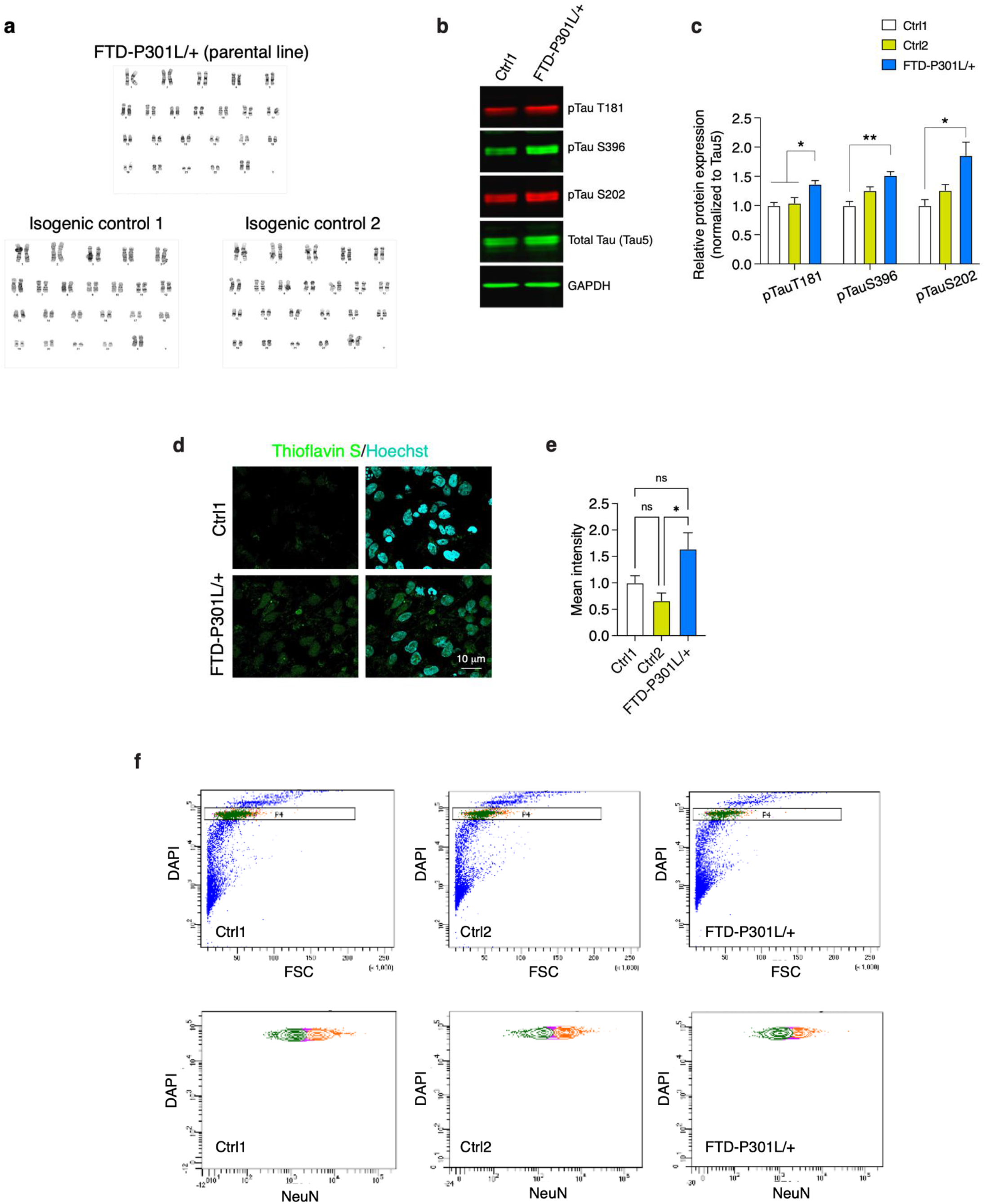
Characterization of human cerebral organoids derived from FTD patient iPSC line and isogenic control lines. **a** FTD patient iPSC line and two isogenic control lines exhibit normal karyotype. **b** Representative blot images of pTau (T181, S396, S202), Tau5, and GAPDH. **c** Bar graph represents the immunoreactivity for pTau (T181, S396, S202) normalized to isogenic control organoids. **d** Representative images of 60-day-old organoid sections stained for Thioflavin S (green) and Hoechst (cyan). **e** Bar graph represents the intensity of Thioflavin S stain. Three organoids were analyzed in each group. **f** Gating strategy for the isolation of NeuN^+^ and NeuN-nuclei from 3-month-old organoids. All values are shown as mean ± SEM. Statistical analysis: **c, e** one-way ANOVA with Tukey’s post hoc test. Source data are provided as a Source Data file.

**Supplementary Figure 4.**
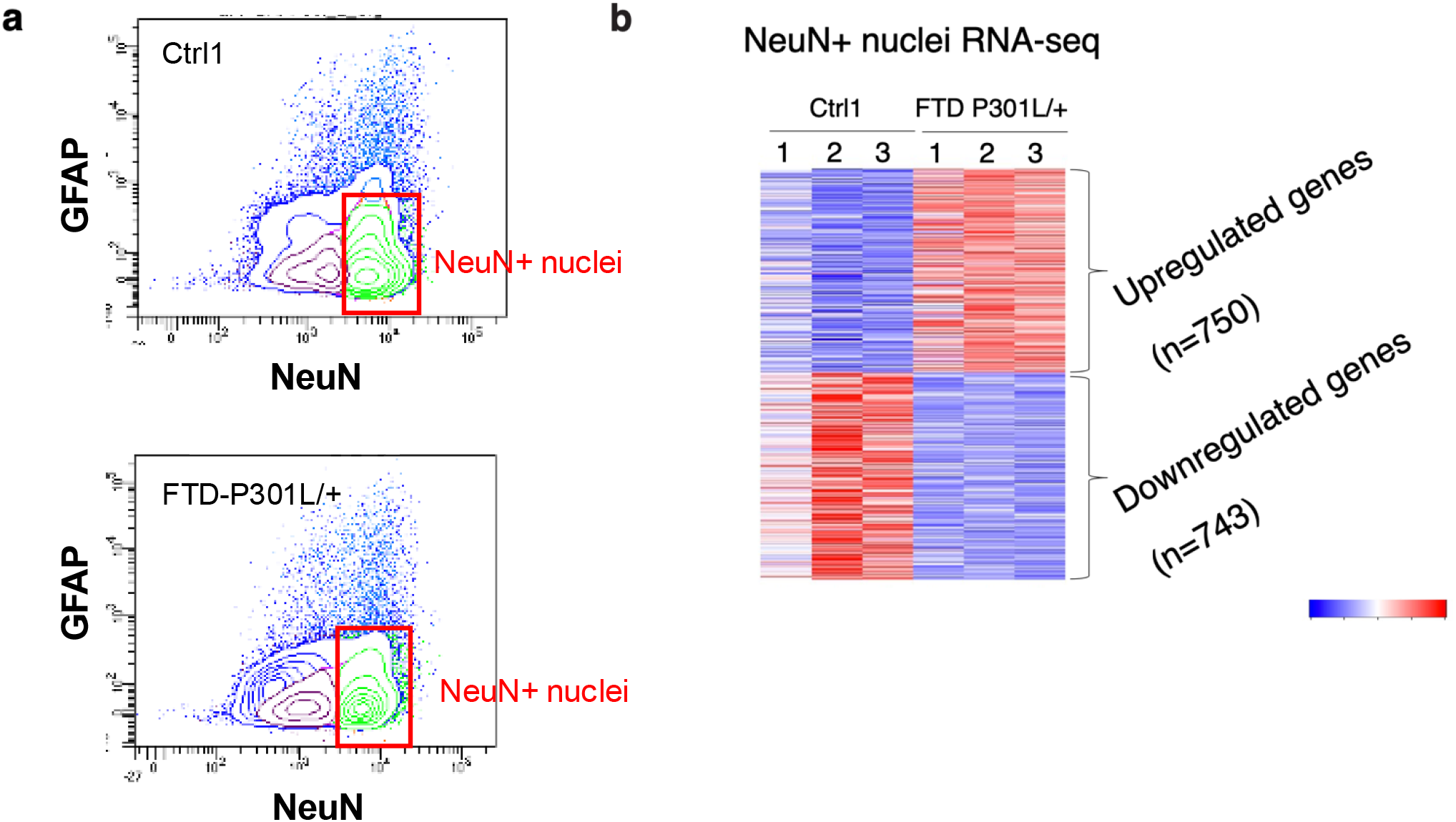
RNA-seq analysis on NeuN^+^ nuclei isolated from FTD human organoids. **a** Gating strategy for the isolation of NeuN^+^ nuclei from 90-day-old organoids. **b** Heatmap of differentially expressed genes (DEGs) that were identified in FTD NeuN^+^ nuclei. Three organoids were analyzed in each group.

## Supplementary Table

**Supplementary Table 1. Differentially expressed gene list in Nutlin-3 treated cultured neurons**

**Supplementary Table 2. Overlapping upregulated genes between Nutlin-3 treated cultured neurons and human AD neurons enriched with NFTs**

**Supplementary Table 3. Differentially expressed gene list in FTD-NeuN**^**+**^ **nuclei; overlapping upregulated genes between Nutlin-3 treated cultured neurons and FTD-NeuN**^**+**^ **nuclei**

## Methods

### Cell culture, treatment, viral transduction, MEA recording, and cell viability assay

Primary cortical neurons were cultured from Swiss–Webster embryos (embryonic day 16), and then were maintained in neurobasal media (Gibco, 21103) supplemented with L-glutamine (5 mM), penicillin and streptomycin, and B27 neuronal additive. Cells were maintained at standard environmental conditions (97% humidity, 5% CO2, 37°C). At 15 DIV, wild-type primary cortical neurons were treated with different compounds for 2 days, including Lig4 inhibitor (SCR-7, 5 µM), p53 inhibitor (Pifithrin-α, 10 nM), p53 activator (Nutlin-3, 100 nM). For I-PpoI-ER experiments, wild-type primary cortical neurons at 5 DIV were transduced with lentivirus expressing I-PpoI-ER-RFP as previously described(Dobbin *et al*, 2013). I-PpoI-ER was induced by adding 4-OHT (Sigma; H7904) to a final concentration of 10 nM and incubated for 2 days prior to assessments. The MEA recordings were conducted at 15DIV and 17DIV, the time point before and after perturbations. Each recording is 20-min long. Cell viability was determined by the CellTiter-Glo® Luminescent Cell Viability Assay (Promega).

### iPSCs cultures

Original approval for human subject work to derive the iPSC line was obtained under a Partners/Massachusetts General Hospital-approved Institutional Review Board Protocol (#2010P001611/MGH)(Seo *et al*, 2017). The healthy human control iPSC line (CS-0009-01) was acquired from McClean. The FTD patient (Tau P301L +/-) iPSC line (MGH2046) was used to generate two clonal isogenic control lines with the P301L mutation in *MAPT* was corrected to P301P by CRISPR/Cas9 genome editing. All lines were cultured on irradiated mouse embryonic fibroblasts (MEFs) in DMEM/F12, HEPES media (Gibco) supplemented with 20% knockout serum replacement (KSR) (Gibco), 1X non-essential amino acids (NEAA), 1X GlutaMAX, (Life Technologies, Carlsbad, CA), β-fibroblast growth factor (FGF2, PeproTech), and 0.1 mM 2-mercaptoethanol (Sigma-Aldrich, St. Louis, MO). Cultures were passaged weekly using mechanical methods and quality control measurements including stem cell marker expression and Geimsa band karyotyping were performed prior to organoid generation.

### Generation of isogenic control line

A sgRNA for targeting the P301L site in the *MAPT* gene (AACATTAAGCATGTCCCGGG) was designed using http://crispr.mit.edu. The HDR repair template was designed to correct the pathogenic P301L mutation and introduce 4 silent (synonymous) mutations to improve precise editing efficiency. Both sgRNA and repair template were synthesized by IDT. The CRISPR/Cas9 plasmid (pSpCas9-2A-GFP, PX458) was purchased from Addgene, and the sgRNA was cloned into the plasmid and electroporated into iPSCs, followed by FACS, colony inspection, and Sanger sequencing to determine positive clones, as described previously(Seo *et al*, 2017).

### Cerebral organoid culture and calcium imaging

Cerebral organoids were generated from iPSCs as described previously(Seo *et al*, 2017; Raja *et al*, 2016). Following a published protocol(Qian *et al*, 2016), we employed 3D printing to build mini-bioreactors that support growth of uniform cerebral organoids from iPSCs. In brief, embryoid bodies (EBs) were formed by loading 12,000 iPSCs per well into 96-well plates pre-coated with Pluronic acid (1%, F-127, Sigma-Aldrich) and maintained in media consists of Glasgow-MEM supplement with 20% KSR, 1X sodium pyruvate, 1X NEAA, 0.1 mM 2-mercaptoethanol, 20 μM ROCK inhibitor (Y-27632, Millipore), 5 μM TGFβ-inhibitor (SB431532, Tocris Biosciences, Bristol, UK), and 3 μM Wnt-inhibitor (IWRe1, Tocris Biosciences) for 20 days. Cerebral organoids were then transferred to non-adherent petri-dishes and cultured in media consisted of DMEM/F12 supplemented with 1X Chemically Defined Lipid Concentrate and 1X N2-supplement to promote neuroepithelial formation. From day 35, 5 μM heparin (Sigma-Aldrich), 10% FBS, and 1% matrigel (Life Sciences) were added to the medium. From day 70, 1X B27-supplement and 2% matrigel (Life Sciences) were added to the medium. For calcium imaging, organoids were transduced with pAAV.Syn.GCaMP6s.WPRE.SV40 (Addgene; 100843-AAV1). Two weeks after viral transduction, organoids were treated with different compounds for 7 days, including Lig4 inhibitor (SCR-7, 5 µM), p53 inhibitor (Pifithrin-α, 10 nM), p53 activator (Nutlin-3, 100 nM). For live imaging, each organoid was transferred to a U-shaped 96-well plate and imaged using a Zeiss LSM 710 confocal microscope for 20 minutes. The calcium dynamic was determined using the average ROI. ΔF/F compares the changes in the intensity to the baseline over a 20-minute recording.

### Electrophysiological Recordings

Cells were removed from the incubator, and incubation culture media was replaced with an oxygenated artificial cerebrospinal fluid (ACSF) solution containing (in mM) 124 NaCl, 3.3 KCl, 1.3 NaH2PO4, 2.5 CaCl2•2H2O, 1.5 MgCl2, 26 NaHCO3 and 11 D-Glucose. Cells were placed in a recording chamber and perfused with oxygenated ACSF at a constant rate of 2 ml/min at room temperature (24°C). Cells were visualized using infrared differential interference contrast imaging on an Olympus BX-50WI microscope. Electrophysiological recordings were performed using an Axon Multiclamp 700B patch-clamp amplifier (Molecular Devices) and Clampex software (version 11.2, Molecular Devices). Signals were filtered at 1 kHz using the amplifier’s four-pole, low-pass Bessel filter, digitized at 10 kHz with an Axon Digidata 1550B interface (Molecular Devices) and stored on a personal computer. Spontaneous excitatory postsynaptic current (sEPSC) was acquired in the whole-cell voltage-clamp configuration and gap-free acquisition mode. Cells were held at a membrane potential of −70 mV for sEPSC. Action potentials were generated by injecting currents in the whole-cell current-clamp configuration. Recording electrode solution contained (in mM) 120 K gluconate, 5 KCl, 2 MgCl2•6H2O 10 HEPES, 4 ATP, 2 GTP. sEPSC frequency and amplitude was detected using Mini Analysis Program (Synaptosoft Inc. Fort Lee, NJ). Action potentials were analyzed using a threshold detection protocol of Clampfit software (version 11.2, Molecular Devices).

### RNA-seq analysis of mouse cultured neurons

Total RNA was extracted from mouse cultured neurons using TRIzol reagent (Invitrogen). Purified RNA was subjected to quality control (Fragment Analyzer). cDNA libraries were prepared using NEB ribosomal depletion kit (NEB). Libraries were sequenced on the Illumina Nextseq 75 platform at MIT BioMicro Center. FASTQ paired-end reads were aligned using STAR (v.2.6.1a) to GRCm339 reference genome. Transcripts were quantified using HTSeq, data was normalized using RUV-seq and differential gene expression analysis was performed using DEseq2(Risso *et al*, 2014; Love *et al*, 2014). Differentially expressed genes were called with a *p*.*adj* < 0.05, log_2_FoldChange > 0.2 or < −0.2. Genes with > 10 normalized counts in at least one sample were considered expressed. p53 target genes were extracted from (V$P53_02 and V$P53 DECAMER Q2) lists in the transcription factor binding site (TFBS) database. Gene ontology (GO) analysis of DEGs were performed using Toppgene. Data were deposited in the GEO Data Bank under ID code GSE251858. The lists of differentially expressed genes can be found in **Supplementary Table 1**.

### Single-cell electrophoresis (comet) assay

Hippocampal tissues or single-cell suspensions were homogenized in 0.5 mL cold PBS using a rotor stator on ice. After homogenization, cells were run through a 0.45 μm cell strainer and diluted with cold PBS to 4 × 10^5^ cells/mL. For comet after FACS, freshly isolated NeuN^+^ nuclei were used. Fifty microliters of diluted cells were added to 450 μL of 1% low melting point agarose (Invitrogen) kept at 42 °C and applied onto each well of comet slides (Trevigen, 4250-004-03). Comet slides were kept at 4 °C in the dark for 30 min, and then immersed in cold lysis solution (Trevigen, 4250-050-01) that contained 10% dimethyl sulfoxide (DMSO) at 4 °C in the dark for 14 to 16 h. After overnight lysis, slides were incubated with freshly prepared pre-chilled neutral buffer (0.3M sodium acetate, 0.1M Tris, pH 9.0) for 1 h in the dark at 4 °C and then subjected to electrophoresis in 850 mL cold neutral buffer using CometAssay® Electrophoresis System II (Trevigen, 4250-050-ES) following the manufacturer’s instructions. After electrophoresis, slides were incubated with DNA precipitation solution (6.7mL 7.5M NH4Ac, 43.3mL 95% EtOH) for 30 min at RT, and then incubated with 70% ethanol at RT for 30 min. Slides were next dried at 37 °C for 30 min to bring cells into the sample plane, and stained with SYBR Gold (Invitrogen, S11494) for 30 min at RT. Slides were imaged using a Zeiss LSM710 confocal microscope (×5 objective) with tile scan. Comet images were analyzed using default settings in the OpenComet software, a free plugin for ImageJ (http://www.cometbio.org/index.html)(Pao *et al*, 2019).

### Immunostaining in cultured neurons and organoid section

Organoids were then embedded in optimal cutting temperature compound (OCT compound) and sliced as 20-µm sections using a Cryostat microtome (Leica). For immunostaining,cultured neurons or organoid sections were fixed in 4% formaldehyde for 10 min at room temperature, followed by a PBS wash. Sections or cells were permeabilized and blocked in PBS containing 0.2% TritonX-100 and 10% normal donkey serum for 1 hour at room temperature, then incubated with primary antibody at 4 °C overnight. Primary antibody was visualized using the appropriate secondary antibody conjugated to Alexa Fluor 488, Alexa Fluor 594, or Alexa Fluor 647 (Thermo Fisher Scientific). Nuclei were visualized with Hoechst 33342 (Thermo Fisher Scientific). All images were captured using a Zeiss LSM 710 or 880 confocal microscope and the ZEN software, and analyzed using the ImageJ software (National Institutes of Health, https://imagej.nih.gov/ij/, RRID: SCR_003070). For each experiment, two sections from one organoid or two images from one coverslip were used for quantification.

### Immunostaining in post-mortem human brain sections

Frozen postmortem cortical samples (prefrontal cortex) of three healthy subjects and three subjects with AD pathology were from the Religious Orders Study and Rush Memory and Aging Project (ROSMAP). Brain tissue of approximately 0.1 cm^3^ in volume was immersed in 4% PFA at 4°C overnight before embedded into 4% low-melting point agarose. The embedded brain tissue was then sectioned at 40 µm using a vibratome (Leica), and the sections were post-fixed in 4% PFA overnight at 4°C. For antigen retrieval, the brain sections were incubated at 95°C in antigen retrieval solution (10 mM sodium citrate, 0.05% Tween-20, pH6.0) for 45 min and then placed in PBS for 20 min at room temperature. After washing with PBS, sections were incubated at 37°C in 2N HCl for 30 min followed by washes with 50 mM Tris-HCl pH7.6. The sections were then permeabilized in blocking solution (5% BSA, 0.2% Triton-X100 in PBS) for 1 h at room temperature. The sections were incubated for 72 h at 4°C in primary antibody in blocking solution. Primary antibodies were anti-AT8 (1:250; Thermo Fisher Scientific #MN1020), anti-γH2AX (1:250; abcam ab11174), and anti-NeuN (1:1000; Synaptic Systems #266004). Primary antibody was visualized using the appropriate secondary antibody conjugated to Alexa Fluor 488, Alexa Fluor 594, or Alexa Fluor 647 (Thermo Fisher Scientific). All images were captured using a Zeiss LSM 900 confocal microscope and the ZEN software and analyzed using the Imaris image analysis software (Bitplane). NeuN signals were used to create a mask for measuring the intensity of AT8 and γH2AX within NeuN mask. Four sections from each subject were used for quantification.

### Antibodies

γ-H2AX (05-636, Millipore; ab11174, abcam), pATM S1981 (ab36810, Abcam), pp53 S15 (PA5-64821, Thermo Fischer Scientific), pTau T181 (9632, Cell Signaling Technology), pTau S202 (11834S, Cell Signaling Technology), pTau S396 (5383, Cell Signaling Technology), pTau S404 (20194, Cell Signaling Technology), pTau S202/T205 (MN1020, Thermo Fischer Scientific), GAPDH (2118, Cell Signaling Technology), NeuN (266 004, Synaptic Systems), MAP2 (822501, BioLegend). Synaptophysin (MA1-213, Thermo Fisher Scientific), CTIP2 (ab18465, Abcam), TBR1 (ab183032, Abcam).

### RNA-seq analysis of NeuN^+^ nuclei from cerebral organoids

FACS isolated NeuN^+^ nuclei were immediately used for RNA-seq library generation. RNA was extracted from nuclei using the RecoverAll Total Nuclei Acid Isolation Kit for FFPE Tissue (ThermoFisher). The amount and the quality of purified RNA were determined by the Femto Pulse Systems (Agilent, Santa Clara, CA). RNA libraries were constructed using the Low Input Mammalian Total Clontech ZapR Kit (Clontech, Mountain View, CA) and sequenced using a Nextseq500 platform (Illumina, San Diego, CA) with 40 bp paired-end reads (40PE). Sequencing was performed at the MIT BioMicroCenter (https://openwetware.org/wiki/BioMicroCenter). RNAseq reads were aligned using STAR (version 2.5.4a). The raw fastq reads were aligned by Tophat2, and further processed by Cufflinks 2.0.0 with UCSC hg19 reference gene annotation to determine transcript abundances. A gene was considered differentially expressed with a statistical significance of *adj P* < 0.05. Gene ontology (GO) analysis of DEGs was performed using Toppgene. The lists of differentially expressed genes can be found in **Supplementary Table 3**.

### Statistics

Results are shown as mean ± SEM. All statistical analysis was conducted using the Prism GraphPad software. Two-tailed unpaired t test, two-tailed Fisher’s exact test, one-way analysis of variance (ANOVA), followed by Tukey’s post hoc test, two-way ANOVA followed by Bonferroni post hoc test, and two-way repeated-measures ANOVA followed by Bonferroni post hoc test were used. The data were assumed to be distributed normally, but this was not formally tested. No statistical method was used to determine sample sizes, but sample sizes in this study were similar to previous publications(Dobbin *et al*, 2013; Seo *et al*, 2017). Quantifications of immunoreactivity were performed blind to the conditions of the experiment and were analyzed with the Imaris image analysis software (Bitplane) and ImageJ.

## Contributions

P.-C.P. and L.-H.T. designed the study, and L.-H.T. directed and coordinated the study. P.-C.P. initiated, planned, and performed experiments using mouse cultured neurons, human organoids, and *post-mortem* brain section. L.A.W conducted CRISPR/Cas9 editing on the FTD iPSC line. L.A.W characterized the pathology in FTD organoids. P.-C.P. assessed neuronal firing in human organoids. L.A.W. and F.G. performed RNAseq analysis on FTD organoids. L.W. conducted patch-clamp electrophysiological characterization. L.A.W. performed RNAseq analysis on mouse cultured neurons. A.L. and S.R. contributed to quantification of experiments. A.L. and A.S. prepared and maintained primary neuronal culture. P.-C.P., D.D., and A.L. performed immunostaining on mouse cultured neurons, human organoids, and *post-mortem* brain section. A.L., C.S. and O.K. maintained FTD organoids, and A.L. contributed to the establishment of bioreactor for organoid culture. P.-C.P., U.G., J.P., and L.-H.T. wrote the manuscript with critical input from all the authors.

## Acknowledgements

We thank E. McNamara for mouse colony maintenance. We thank BioMicroCenter and Kock Institute Flow Cytometry core at MIT for their help and guidance on RNA-seq and sorting experiments. We thank the individuals who donated post-mortem brain samples, and their families, for enabling this research. We thank members of Tsai lab for discussion and valuable comments on manuscript. This work was supported by NINDS Grant (NS102730) and Glenn Foundation Award for research in biological mechanisms of aging to L.-H.T.

## Data availability

All data are available from the corresponding author upon reasonable request. The source data are provided as a Source Data file. RNA-seq data is publicly available and can be downloaded at the NCBI Gene Expression Omnibus (GSE251858).

